# Effects of DNA oxidation on the evolution of genomes

**DOI:** 10.1101/150425

**Authors:** Michael Sheinman, Rutger Hermsen

## Abstract

Oxidation of DNA increases its mutation rate, causing otherwise rare G → T transversions during DNA replication. Here we use a comparative genomic approach to assess the importance of DNA oxidation for the evolution of genomic sequences. To do so, we study the mutational spectrum of G*_n_*-tracks on various timescales, ranging from one human generation to the divergence between primates, and compare it to the properties of guanines oxidation known from experimental and computational studies. Our results suggest that, in short G*_n_* tracks (*n* ≤ 3), oxidation does not dominate the mutagenesis of guanines, except in cancerous tumors, especially in lungs. However, we consistently find that the G → T transversion rate is elevated by an order of magnitude in long G*_n_* tracks (*n* ≳ 6). In such long G*_n_*-tracks, G → T substitutions in fact dominate the mutational spectrum, suggesting that long G*_n_* tracks are oxidized more frequently and/or repaired less efficiently.

## I. INTRODUCTION

Oxidative stress has always accompanied life during its evolution. It can damage all components of the cell, with severe consequences. Indeed, oxidation is thought to play a role in the development of many human conditions, including Alzheimer’s disease [1, 2], cancer [3, 4], heart failure [5], and aging [6, 7]. Of particular significance is the oxidation of nucleic acids, which can induce several kinds of covalent modifications to DNA and RNA, such as single-nucleobase lesions, strand breaks, inter- and intrastrand cross-links, and protein-DNA cross-links [8]. Such lesions can cause mutations [9] and form road blocks for transcription [10] and translation [11, 12]. Much effort has been made to understand precisely how nucleotides are oxidized and how this in turn affects transcription, translation, DNA replication and gene expression [13]. Nevertheless, many aspects are still poorly understood. In particular, it is largely unknown how oxidative lesions affect the *evolution* of genomic sequences. Because oxidation causes particular mutations in particular sequences, we ask: Can we identify footprints of oxidative stress in the evolution of DNA sequences of humans and other primates?

As we will detail below, many studies have shown that oxidation particularly affects guanines and tandem repeats of them, here called G*_n_* tracks, where *n* denotes the length of the repeat. In such sequences, oxidation is known to cause otherwise rare G → T substitutions. In this study, we therefore use a comparative genomics approach to analyze the mutational spectrum of G*_n_* tracks. We report several intriguing patterns. Most importantly, while G → T substitutions are rare in short G*_n_* tracks, in long G*_n_* tracks (*n* ≳ 6) they are an order of magnitude more frequent and in fact dominate the mutational spectrum.

Before presenting these results, we first briefly review several relevant properties of DNA oxidation, the associated mutational spectra, and their sequence dependence.

### A. DNA oxidation: three patterns

The *in vivo* oxidation rates of nucleotides vary strongly between organisms [14]. In addition, they depend in nontrivial ways on the particular oxidant: even qualitative patterns can differ between oxidants [15, 16]. Nevertheless, three generic trends have been observed *in vivo, in vitro* and *in silico.*

First, oxidation preferentially affects guanines. *In vitro* studies indicate that, of all DNA bases, guanine is oxidized most readily due to its low redox potential [17]; many oxidants oxidize it selectively [18]. Correspondingly, single strands of poly(G) are oxidized much more readily than poly(A), poly(C), poly(U) and heterogeneous sequences [19].

Second, the oxidation rate per base of a G*_n_* track increases with its length *n,* as has been measured *in vitro* for short sequences [20, 21]. Accumulating computational studies and *in vitro* experiments explain this phenomenon: stacking interactions between adjacent guanines affect their electron orbitals, raising the highest occupied molecular orbital and thus lowering the ionization potential. Indeed, of all 10 possible nucleotide duplets, the G_2_ duplet possesses the lowest ionization potential [16, 22–24]; in addition, the ionization potential of G_3_ is lower than that of G_2_ and the ionization potential of G_4_ is lower than that of G_3_ [22].

Third, the 5’ end of a G*_n_* track has a higher oxidation rate than its 3’ end. Indeed, the highest occupied molecular orbital in stacked guanines is localized at the 5’ end of the G_n_ track. This makes the 5’ end of adjacent guanines an effective sink of electron holes transferring along the DNA molecule [25]. Thus, it is predicted that in G_2_ doublets, the oxidation rate of the 5’ base is an order of magnitude higher than that of the 3’ base [20, 21, 25, 26]. Similarly, in G_3_ triplets the oxidized G occurs preferentially at the first or second position [6, 20, 21].

These three patterns motivate the focus of this study on the evolution of G_*n*_ tracks.

### B. Mutations caused by oxidation: G → T transversions

The lesions resulting from oxidation of guanines can result in mutations. The most common lesion is 8-oxoguanine (8-oxo-G). During DNA replication, 8-oxo-G can pair with adenine in the complementary strand, which results in G → T transversions [27]. Importantly, transversions are otherwise rare: in many organisms, a strong, several-fold prevalence of transitions over transversions has been observed [28] with few exceptions [29, 30]. Abundant G → T transversions are therefore indicative of oxidative stress. For this reason, one would expect the three patterns of oxidation described above to be reflected in the rate of G → T transversions occurring in G*_n_* tracks. Some observations support this expectation; for instance, in *E. coli* the 5’-end guanines of the G_2_ duplets were found to be mutational hotspots [31], predominantly generating G → T transversions [32].

However, the properties of mutational spectra are nontrivial. The mutability of genomic DNA depends on many global factors, such as UV exposure, the concentration of (anti)oxidants and the properties of the error-correction system of the particular organism. In addition, the mutation rate of a nucleotide depends on its local sequence context [33–37]. For instance, CG di-nucleotides in the human genome mutate an order of magnitude faster than other dinucleotides, mostly via G → A and C → T transitions [38]. Furthermore, the overall mutation rate of G·C base pairs is about twice higher than the mutation rate of A·T base pairs [39]. (The high mutability of CG di-nucleotides does not fully account for this difference.) Finally, the context-dependent mutational spectrum can be also affected by molecular details of DNA repair mechanisms and/or the DNA replication process. For instance, in Foster et al. [40] it is suggested that DNA polymerase incorporates A opposite to 8-oxo-G in the template strand more readily if it has just inserted a G opposite a template C.

For the current study, it is particularly important that sequences consisting of tandem repeats tend to mutate faster than generic, high complexity sequences [41]—after all, G_n_ tracks are tandem repeats. Indeed, in several organisms, including *E. coli* [42], *S. cerevisiae* [43] and cultured mammalian cells [44], G*_n_*-tracks are especially prone to various types of mutations, even compared to other low-complexity sequences. While oxidative lesions could contribute to this phenomenon, so do other mutational pathways.

### C. Origins and functions of G*_n_* tracks

Despite their high mutability [44], long G*_n_* tracks are enriched (relative to randomized sequences) in non-RNA-coding parts of large genomes [45, 46]. The origin or function of this enrichment is unclear. Certainly, in most organisms G*_n_* tracks of length *n* = 3 play a role in telomeres [47]. Oxidation preferentially damages these telomeric G*_n_* tracks [48, 49], which induces single-strand breaks and thus contributes to telomere shortening [50]. Furthermore, the enrichment of G*_n_* tracks could be related to the ability of certain G-rich DNA sequences to form G quadruplex DNA, a non-canonical DNA structure [51]. Such structures are found to be related to neurodegenerative diseases [52] and cancer [53, 54]. Lastly, it has been suggested that G*_n_* tracks in genomes may act as traps of electrons transferring across the base stack [23], thus protecting important parts of a genome from lesions and mutations [55–58]. This squares well with the intriguing observation that intronic regions close to exons are enriched with G*_n_* tracks [57–59], but direct evidence for such a protection is still lacking [60].

In contrast, in RNA-coding sequences long G*_n_* tracks are avoided [61]. A possible explanation is that transcription of oxidized DNA is problematic [62]. Incidentally, this would also explain certain patterns observed in RNA-seq data. For instance, in the blood RNA-seq of patients with coronary artery calcification—a condition related to oxidative stress [63]—G_6_ and C_6_ are the only two significantly depleted 6-mers [64].

### D. Repair mechanisms

In all kingdoms of life, enzymes have been identified that excise 8-oxo-G, repair it, or control its damage [65]. Mutations in such repair enzymes affect mutational patterns in consistent ways.

In *E. coli,* the repair enzymes are called MutY, MutM, and MutT. In mutation strains unable to prevent or repair oxidative DNA damage, the rate of G → T transversions is significantly increased, making them the dominant substitution process for guanines [40, 66–69]. In *S. cerevisiae,* the Ogg1 protein is a functional analog of the bacterial MutY (even though these enzymes do not show significant sequence homology [70]). Ogg1-deficient strains specifically accumulate G → T transversions [71, 72]. In humans, the mammalian homolog of the *S. cerevisiae* OGG1 gene, called hOGG, is the primary enzyme responsible for the excision of 8-oxo-G [66]. Mutations in this enzyme are associated with an elevated risk for various cancers [73–77] and its reduced expression is associated with Alzheimer disease [78]. *C. elegans* is the only known organism without any 8-oxo-G repair component homologous to either those of *E. coli* or yeast [65]. In this light, it is striking that in mutation-accumulating *C. elegans* lines, G → T transversions dominate over G → A transitions [30].

To sum up, we expect that the mutational spectra of G*_n_* tracks observed in chromosomes of living organisms should be affected by the combined effect of sequence-dependent oxidation rates, damage-repair mechanisms, mutational pathways, and selection. Below, we analyze genomic sequences of humans and other primates to assess the role of DNA oxidation in the evolution of the genome sequences on different timescales. Using G → T transversions to estimate the rate of unrepaired oxidative lesions, we ask: Can the before-mentioned properties of guanine oxidation be observed in the mutational patterns of DNA?

## II. RESULTS

### A. The evolution of short G tracks

We start by analyzing the mutational patterns of short G*_n_* tracks of up to three Gs. We begin with arguably the most direct evidence: *de novo* mutations.

#### 1. De novo *substitutions in human parents-children trios*

We combine two datasets [39, 79] containing 15, 953 *de novo* point mutations found by sequencing 328 human parent–offspring trios (see Materials and Methods Section). Fig. 1A shows the substitution probability for guanines within G*_n_* tracks of length *n* = 1, 2 or 3 *(i.e.,* HGH, HGGH and HGGGH sequences, where H denotes a non-G nucleotide). The stacked bars display the distribution over the possible substitutions: G → T (red), G → C (blue), or G → A (green).

**Figure 1:**
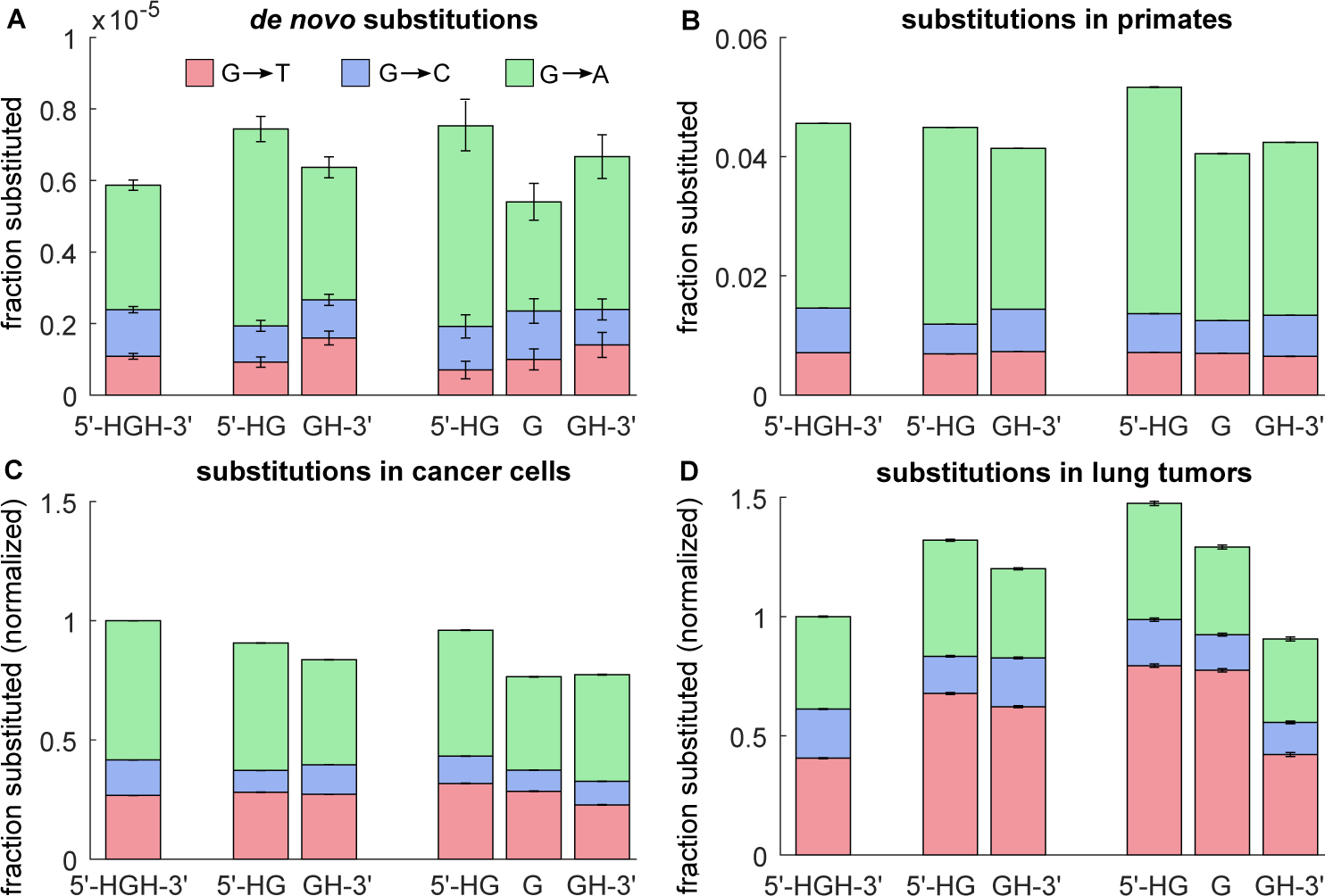
Mutational spectra of guanines in at each position of short G*_n_* tracks (HGH, HGGH, and HGGGH sequences), based on four data sets. Error bars represent 95% confidence intervals. (*A*) Fraction of Gs mutated to T, C and A in a single generation, derived from sequenced parent–offspring trios [39, 79]. (*B*) Fraction of Gs mutated to T, C, and A in 7 primates, relative to their reconstructed common ancestor [80, 81]. (C) Estimated rate of somatic substitutions in tumorous cells, based on the COSMIC database [82]. All rates are normalized to the substitution rate of HGH guanines (first bar). (D) Same as panel *C,* but for the subset of lung tumors.

Even though the mutation rates at the various positions are similar, several significant patterns are found. Overall, G → A substitutions dominate the mutational spectrum in short G*_n_* tracks, accounting for (63 ± 1)% of the substitutions. In G_2_ di-nucleotides, G → A substitutions occur more frequently at the 5’ G than at the 3’ G; this pattern can be explained by the high rate of G → A mutations in CG di-nucleotides [38].

In these short tracks, G → T transversions make up (18±1)% of the substitutions. Based on the properties of DNA oxidation reviewed in the introduction, we expected that the G → T rate per guanine would increase with the length the G*_n_* track. This is not borne out by these data: the rate per guanine is similar and near 1 × 10^-6^ for tracks of lengths 1 to 3. In addition, we expected that G → T substitutions would be biased to the 5’ end of the tracks. Surprisingly, the opposite is true: in G_2_ sequences, 5’ Gs mutate (42 ± 8)% *less* frequently than 3’ Gs do (binomial test, *p* = 2 × 10^-8^). In G_3_ tracks, the G → T rate also seems to increase from 5’ to 3’—but here the larger error bars do not permit firm conclusions (χ^2^ = 10.6, df = 2, *p* = 0.01).

#### 2. Substitutions in primates

We now ask whether similar trends are also visible in the mutations that have accumulated for tens of millions of years during the evolution of the primates [83]. To obtain the mutational spectrum of guanines on this timescale, we analyze a multiple alignment of 7 primate genomes, from *Homo sapiens, Pan troglodytes, Gorilla gorilla, Pongo abelii, Papio anubis, Callithrix jacchus,* and *Macaca mulatta* [80, 81]. We identified substitutions in short G*_n_* tracks by comparing each genome to the reconstructed common ancestor (see Materials and Methods Section).

The results are summarized in Fig. 1B. Overall, the substitution frequencies found here are very similar to those observed in the parent–offspring trios (Fig. 1A). Again, G → A substitutions make up the bulk of the substitutions (69%) and are biased towards the 5’ end of the G*_n_* track due to the high mutability of CG di-nucleotides. Similar as before, G → T transversions account for 16% of all substitutions. In G_2_ sequences, G → T transversion are again more frequent in 3’ Gs, even though the difference of (5.8 ± 0.2)% is clearly smaller than in the *de novo* mutations. But, unexpectedly, in G_3_ tracks the trend is reversed: the G → T transversion rate now decreases from 5’ to 3’ (x^2^ = 10^6^, df = 2, *p* < 10^-100^), more in line with the expectation based on the computational and *in vitro* studies. We note, however, that the effect is small in an absolute sense.

#### 3. Somatic mutations in cancer

Because cancer is frequently associated with oxidative stress, one would expect an elevated frequency of G → T transversions in cancer cells. We analyzed the somatic mutations included in the Catalogue Of Somatic Mutations In Cancer (COSMIC) dataset [82]; the mutational spectrum of short G*_n_* tracks based on the full database is shown in Fig. 1C.

Indeed, G → T transversions are much more frequent in the cancer data: they make up 29% of the substitutions, compared to 18% in the parent–offspring trios. Also, we now find a clear preference for G → T substitutions to occur at the 5’ end of the tracks. In G_2_ tracks, the effect is very small but statistically significant (binomial test, *p* < 10^−8^). In G_3_ tracks, the effect is clearer (χ^2^ = 10^6^, *p* < 10^−100^): the 5’ G is mutated (40 ± 1)% more frequently than the 3’ G. Both are in stark contrast to the opposite trend in the *de novo* mutations in parent–offspring trios.

The COSMIC dataset contains mutations for many cancer types, which are not equally associated with oxidation. Smoking-associated cancers are particularly associated with oxidation. This is reflected in mutation patterns. For instance, it is known that cancer cells in smoking-associated tumors exhibit a higher G → T transversion rate [84]. Also, the cancer signature associated with smoking is the only one in which the G3 triplet has the highest mutation rate of all triplets [85]. This signature is similar to the mutational pattern observed in experimental systems exposed to tobacco carcinogens [85]. We therefore repeated the analysis on a subset of the COSMIC database containing mutations in lung tumors only.

The results are, for the first time, fully in line with our expectations (see Fig. 1D). In lung tumors, (46 f 0.1)% of the substitutions in short G*_n_* tracks are G → T transversions—2.6 times more than in the *de novo* mutations. Guanines in G_2_ tracks now mutate (60 f 1)% faster than G_1_ sequences do. Moreover, guanines in the 5’ or middle position of G_3_ tracks mutate even more frequently. Lastly, in G_3_ tracks, there is an obvious trend of a higher G → T frequency towards the 5’ end: the G → T rate of the 5’ G is (86 ± 2)% larger than that of the 3’ G.

To summarize, the above analyses yield the following picture. The *de novo* mutations found in parent– offspring trios, and those inferred from the primate lineages, did not confirm the expectations based on computational and *in vitro* studies of oxidation. In particular, the observed preference of G → T substitutions for the 3’ ends of G*_n_* tracks directly contradicts the prediction. However, in cancer cells, where high levels of oxidative stress are the norm, the trend is reversed. This is particularly striking in lung cancers, where we additionally found that the rate of G → T substitutions increases with the length of the G*_n_* track, as predicted.

### B. The evolution of long G tracks

Although *in vitro* data and computational predictions are lacking for oxidation rates of G*_n_* tracks longer than *n* = 3, we now extend our analysis to the substitution spectrum of long G*_n_* tracks.

#### 1. De novo *substitutions in human parents-offspring trios*

We again start with *de novo* substitutions in human parent–offspring trios, using the same datasets as before [39, 79] (see Materials and Methods). Because long G*_n_* tracks are rare and the dataset is small, no substitutions are found in G*_n_* tracks with a length *n >* 6. Shorter tracks, however, do show a clear trend.

In Fig. 2A the overall substitution rate per nucleotide is shown as a function of the G-track length *n.* It appears that long G*_n_* tracks are more prone to substitutions than short ones; overall, guanines in G_6_ tracks mutates about twice faster than those in G_1_ tracks.

**Figure 2:**
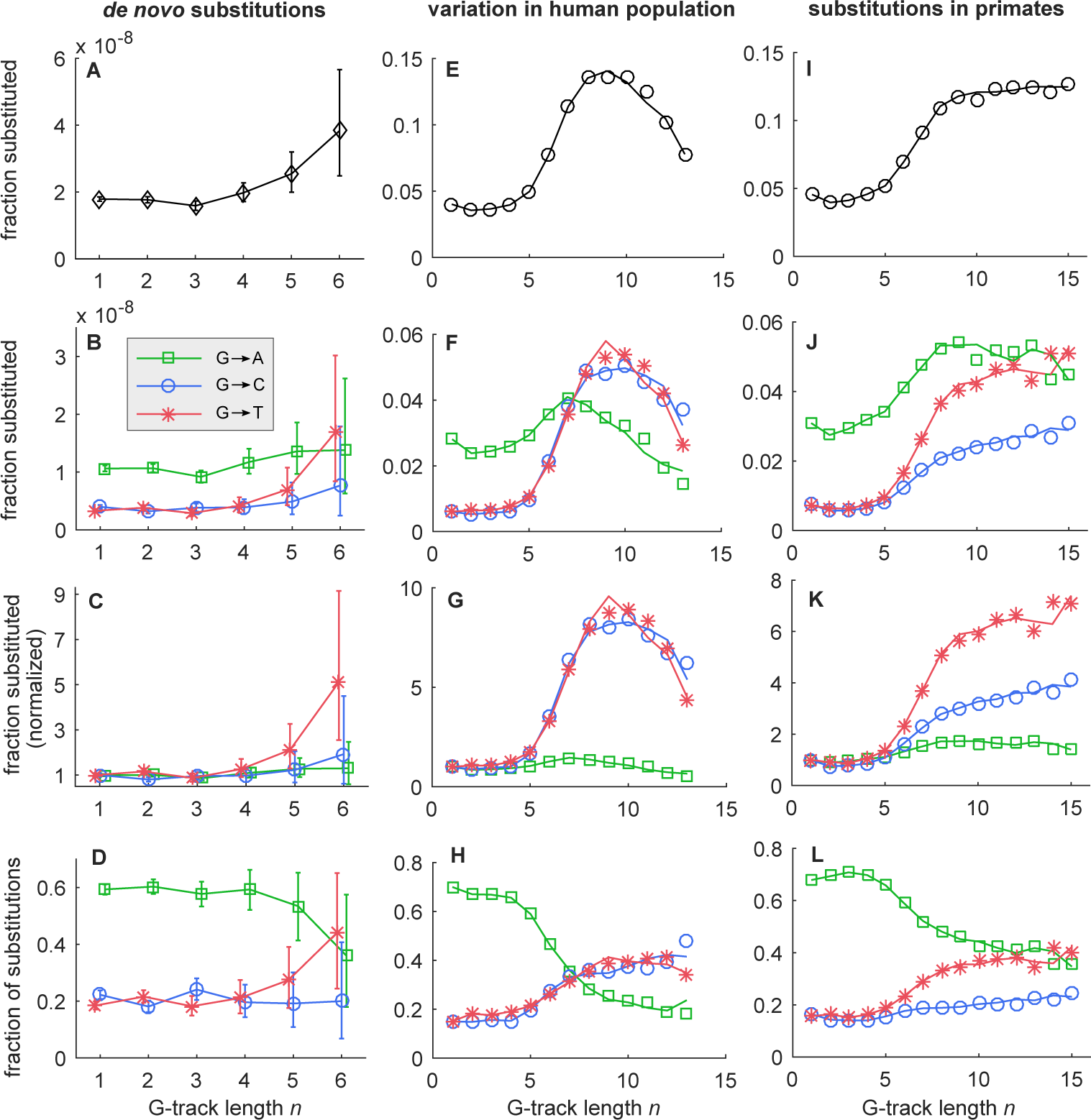
Substitution frequencies per nucleotide in G*_n_* tracks as a function of their length *n,* for three data sets. (*A*–*D*) *De novo* substitution frequencies estimated from DNA sequences of human parent–offspring trios. (*A*) The overall substitution rate per bp increases two-fold as *n* increases from 1 to 6. (*B*) Rates of G → A, G → C and G → T substitutions. The G → T rate increases sharply with *n.* (*C*) Same as (*B*), but now the rates are normalized to 1 at *n* = 1, showing a 5-fold increase in the G → T rate. (*D*) Proportions of the G → A, G → C and G → T substitutions versus *n.* At *n* = 6, G → A transitions no longer dominate the mutational spectrum. (E–H) SNPs of G nucleotides along G*_n_* tracks of length *n* in the human population (data: 1000G project [86]). The symbols and lines present data derived from the plus and minus strands, respectively. (*E*) The proportion of nucleotides in a G*_n_* track of the reconstructed ancestral genome that are identified as SNPs by the 1000G data, as a function of *n.* In line with the *de novo* data (*A*–*D*), an ≈ 6-fold increase in variability is seen as *n* increases above 6. (*F*) Same as (*E*), but now broken down by the alternative alleles A, C, and T (see legend in panel *B*). (*G*) Same as (*F*), but now the rates are normalized to 1 at *n* = 1. A ≈ 6-fold increase in the G → T rate and G → C rate is observed. (*H*) The relative frequencies of the three alternative alleles A, C, and T as a function of *n.* (*I*–*L*) Substitutions along G*_n_* tracks of length *n* in 7 primates, relative to their reconstructed common ancestor. The symbols and lines present data derived from the plus and minus strands, respectively. (*I*) The overall substitution rate per bp as a function of *n* increases more than two-fold as *n* increases above 6. (*J*) Rates of G → A, G → C and G → T substitutions (see legend in panel *B*). While all mutation rates increase with *n,* the G → T and G → C transversion rates have a much larger fold-change than the G → A transition rate. (*K*) Same as (*J*), but now the rates are normalized to 1 at *n* = 1, showing an ≈ 9-fold increase in the G → T mutation rate. (*L*) Proportions of the G → A, G → C and G → T substitutions versus *n.* In long G*_n_* tracks, G → A transitions and G → T transversions have similar frequencies.

In Fig. 2B the total mutation rate is broken down by the alternative substitutions (A, C, or T). The data suggest that they behave differently as a function of *n:* whereas the G → A and G → C substitution rates remain rather constant as *n* increases from 1 to 6, the G → T substitution rate increases about 5-fold. This can be seen most clearly in Fig. 2C, where the fold-change is plotted (*i.e.*, the rate at *n* = 1 is normalized to 1). Consequently, while G → A transitions dominate for the shortest G*_n_* tracks, this is no longer the case at *n* = 6, were G → T transversions are equally prevalent (see Fig. 2D, which shows the relative frequencies of each of the substitutions as a function of n). We note that the error bars of the data grow rapidly with *n* and indicate significant uncertainty at *n* = 6. However, below we will show that the other data sets confirm these trends with much greater statistical confidence.

#### 2. Variability of G_n_ tracks in the human population

The parent–offspring data provide direct evidence for an increasing G → T substitution rate with the length of the G*_n_* track, but they do not provide information on G*_n_* tracks longer than *n* = 6. Therefore, we also analyzed the variability in G*_n_* tracks in the human population. For this, we using data from the 1000G project (Phase 3 release, Sudmant et al. [86]), which identified more than 81 million single-nucleotide polymorphisms (SNPs) based on 2504 individuals from 26 populations.

It is not straightforward to estimate mutation rates from SNP data. We therefore have to make a number of simplifying assumptions. First, we include in the analysis below only the loci with a reconstructed ancestral sequence (reducing the number of SNPs to about 78 million), and defined *n* as the length of the G*_n_* track in the ancestral sequence. Second, because mutations are rare, we assume that each SNP found in the population sample is the result of a single mutation event, even if the same SNP is found in many individuals. (In practice, this means that we ignore the *frequencies* of alleles in the sample.)

The results are shown in Figs. 2(E–H). In these and subsequent figures, we plot the statistics for the two DNA strands separately: the forward strand is represented by symbols, the reverse strand by lines. The fact that the results of both strands are consistent proves that the analysis is free of possible biases or artifacts related to an asymmetric treatment of the strands.

Fig. 2E presents the SNP density (that is, the *fraction* of the Gs that have an alternative allele in the SNP data) in G*_n_* tracks as a function of their length *n.* For lengths *n* ≤ 6, the trend is very similar to that of the *de novo* mutations presented in Fig. 2A–D. (The slightly elevated substitution rate of G_1_ tracks is again explained by the fraction of Gs at the 5’ end that are part of a CG di-nucleotide.) However, as *n* is increased above *n* = 6, the SNP density continues to increase rapidly.

In Fig. 2F the SNP density is split up by the alternative substitutions. While the transition probability of G → A does not depend strongly on *n,* the transversion probability grows ≈ 9-fold (best seen in panel 2G). For short G*_n_* tracks, the alternative allele is typically an A, in line with the general bias of substitutions towards transitions and against transversions. In G*_n_* tracks with *n* > 6, however, transversions dominate the SNP spectrum (see Fig. 2H).

As mentioned above, computational studies suggest that the oxidation rate of G*_n_* tracks is non-uniform along the track [25]. We therefore used the 1000G data to analyze how the SNP frequency depends on the position inside the track. The results are shown in Fig. 3 for *n* = 1 (top row) to *n* = 10 (bottom row). The first column represents the variability of each position, calculated as the fraction of nucleotides at that position that are different from the ancestral genome in at least one individual in the database. To put the variability inside the G*_n_* tracks in context, the figures in this column include the sequences flanking the G*_n_* track; the track itself is indicated with a gray shading. The figures clearly demonstrate that the G*_n_* tracks are much more variable than their flanking sequences. (However, please note the vertical offset of the plots.) More surprisingly, they show that the nucleotides at the borders of the tracks tend to be more variable than the nucleotides in their interior, and that the 5’ ends are generally more variable than the 3’ ends. Columns 2 to 4 of Fig. 3 show the frequencies of the alternative alleles A, C and T as a function of the position along the track. They show that the preference of substitutions for the 5’ ends of long G*_n_* tracks is explained by the strong increase in the frequency of transversions (G → C and G → T) with *n,* which in long G*_n_* tracks (*n* > 6) show a clear preference for the 5’ end.

**Figure 3:**
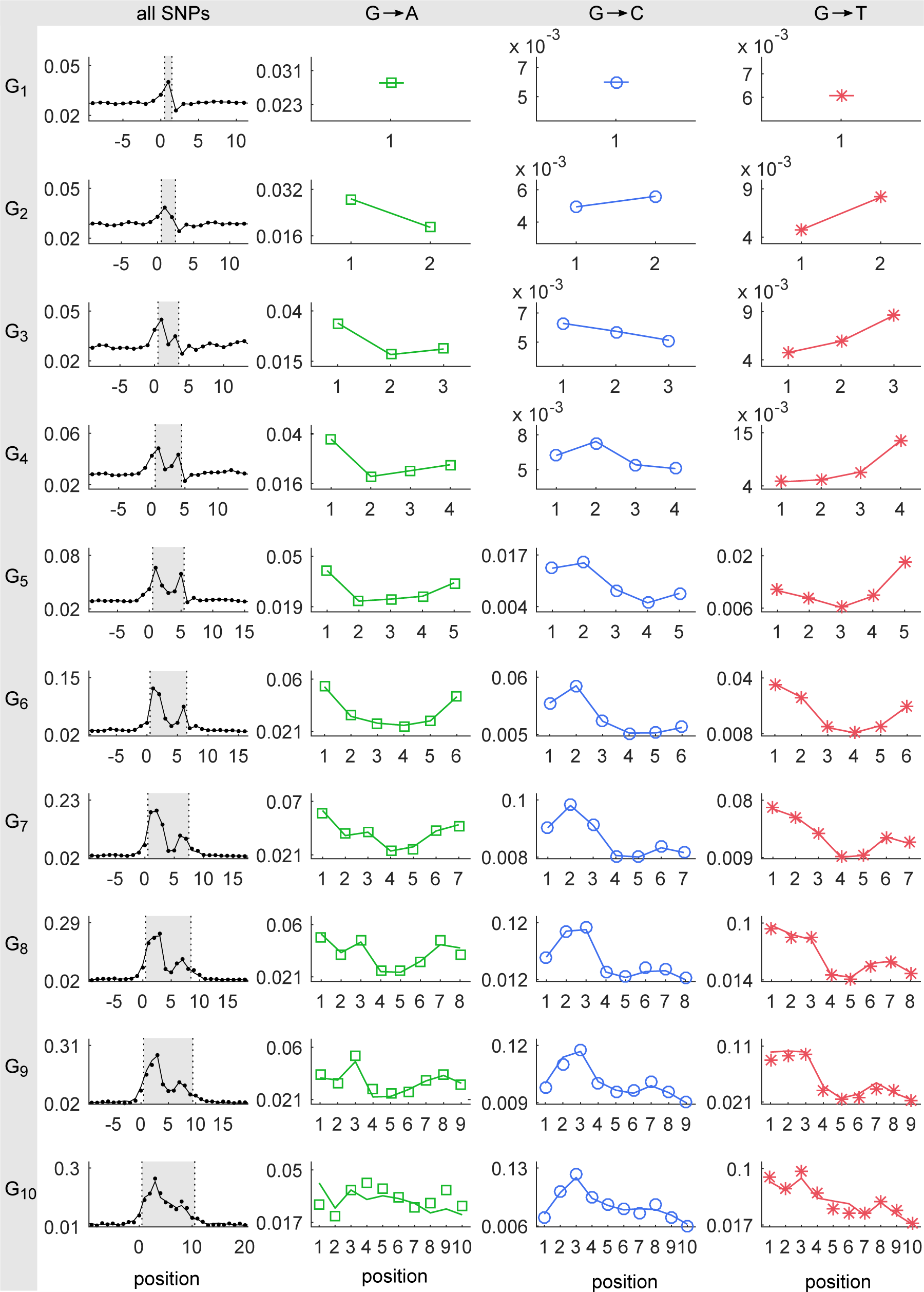
Distribution of SNPs along G*_n_* tracks of the human genome. All G*_n_* tracks of a given length (*n* = 1, 2, …, 10, from top to bottom) in the reconstructed human ancestor were identified and the frequency of SNPs was calculated for each position 1 < *k* < *n* along these tracks. From left to right the first column represents the average fraction of nucleotides that are different from the ancestry genome. The flanks of the G*_n_* track have been included; the G*_n_* tracks themselves are highlighted by the shaded area. The next three columns show the frequencies of SNPs with alternative allele A, C, and T. In all figures, the symbols and lines present data derived from the plus and minus strands, respectively.

#### 3. Substitutions in primates

We now turn to an analysis of the evolution of long G*_n_* tracks in the primate lineages. As for the short G*_n_* tracks, we used the multiple alignment of the genomes of 7 primates and counted substitutions along the G*_n_* tracks by comparing each genome to that of the derived common ancestor.

Fig. 2I plots the substitution rate as a function of *n.* It shows that the substitution rate rapidly increases as *n* exceeds the threshold of *n* ≈ 6, similarly to the trend in human parent–offspring trios and the SNP frequencies in human populations. Fig. 2J demonstrates that, for *n* ≤ 6, the spectrum of substitutions is similar to that of *de novo* substitutions. In long G*_n_* tracks, transversions occur about 6 times more often than in short G*_n_* tracks, while the transition rate exhibits much weaker dependence on *n* (see Fig. 2K). Accordingly, in short G*_n_* tracks substitutions are strongly dominated by G → A transitions, whereas in long G*_n_* tracks this dominance is shared with G → T transversions (Fig. 2K).

Fig. 4 shows the distribution of mutations along G*_n_* as found in the primate lineages. Strikingly, the patterns seen in this figure are highly reminiscent of the distribution of SNPs in the human population data (Fig. 3). Again, the figures in the first column show that G*_n_* tracks are more prone to mutation than their flanking sequences. (Once more, we note that these plots have a non-zero vertical offset.) The nucleotides at the borders of the tracks show a higher mutation rate than the nucleotides in their interior; and in long G*_n_* tracks the 5’ ends are generally more prone to substitutions than the 3’ ends, which is mainly due to a downward slope in the transversion frequencies (blue and red columns).

**Figure 4:**
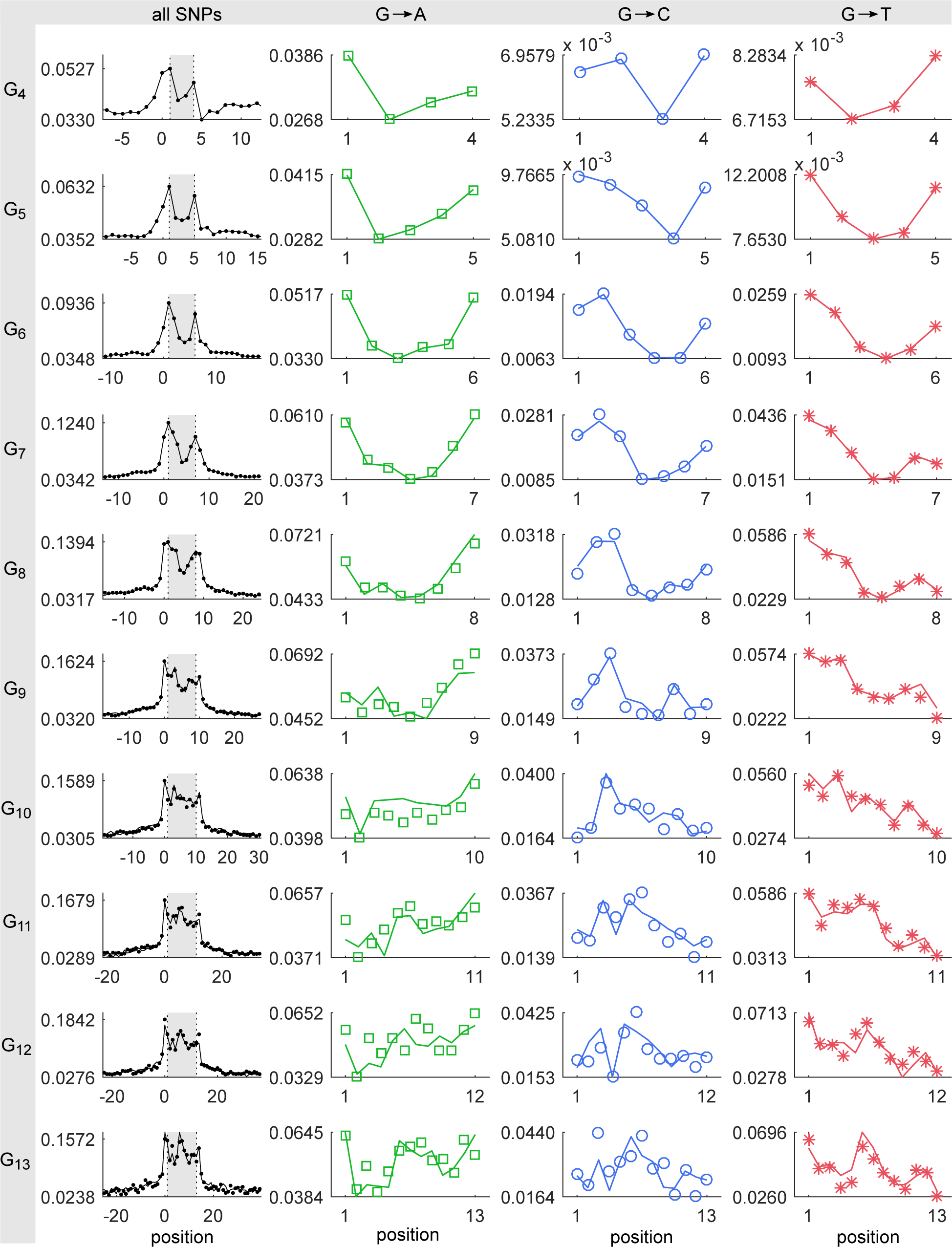
Spectrum of mutations along the G*_n_* tracks of 7 primates lineages, based on alignment of 7 primates and their reconstructed ancestry genome. For a given length of the G*_n_* track (*n* = 4, 5, …, 12, from top to bottom) all G*_n_* tracks were identified in ancestry genome. In the first column the fraction of nucleotides that are different from the ancestry genome is presented. The figures include the flanks of the g tracks; the G*_n_* tracks are highlighted by the shaded area. Columns 2 to 4 show, for each position along the ancestry G*_n_* tracks, the fractions of nucleotides that have been substituted by A, C or T. In all figures, the symbols and lines present data derived from the plus and minus strands, respectively

#### 4. Somatic mutations in cancer

Long G*_n_* tracks are scarce in the human genome. Nevertheless, the COSMIC dataset [82] allows us to estimate the substitution rate of G*_n_* tracks up to *n* = 9. The results are presented in Fig. 5, which shows the relative frequencies of the G → A, G → C, and G → T substitutions as a function of G*_n_*-track length. For long G*_n_* tracks, the G → T transversions dominate among the substitutions. Quantitative interpretation of these data requires caution, because the COSMIC database is a collection of many patients, cancer types, and tissues. Qualitatively, however, the results are in striking accord with the other analyses above.

**Figure 5:**
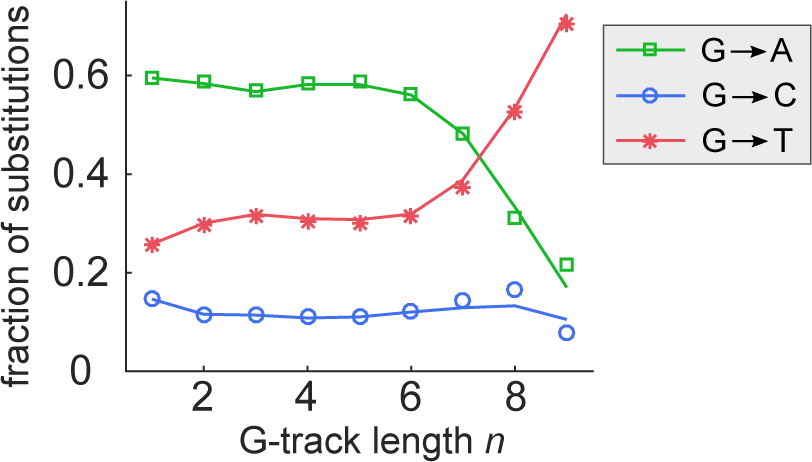
Somatic mutations of G*_n_* tracks in cancerous tumors. Plotted are the relative frequencies of each substitution (G → A, G → C, and G → T) as a function of the track length *n.* The symbols and lines correspond to the plus and minus strands, respectively.

## III. DISCUSSION

In this study, we have analyzed the mutational spectrum of G*_n_* tracks in humans and other primates. Based on the known properties of DNA oxidation and the associated mutations, we expected that the rate of G → T transversions would increase with the length of the track and would show a preference for the 5’ end of the track.

In short G*_n_* tracks (*n* = 1 to 3), our expectations were not borne out. The *de novo* substitutions and the substitutions in the primate lineages show remarkably similar patterns. In neither case does the G → T rate increase appreciably with the length of the G*_n_* tracks; moreover, in the *de novo* substitutions a marked preference is found for the 3’ end rather than the 5’ end of the track. This suggests that the dominant pathway of the G → T transversions in *G_n_* tracks is not related to oxidation. Alternatively, the relevant oxidant could possess a paradoxical oxidation reactivity as a function of ionization potential, as shown by Margolin et al. [15].

The situation is different, however, in cancer cells, and in particular in lung tumors, where the expected trends *are* clearly seen. One possible explanation is that at least two pathways produce G → T substitutions: one independent of oxidation that dominates under normal conditions, and one that does depend on oxidation that dominates under conditions of high oxidative stress and mutagenesis, such as in lung tumors. That said, it is also possible that the different mutational spectrum of short G*_n_* tracks in cancer cells reflects changes in, for instance, their DNA-damage repair machinery.

In long G*_n_* tracks, we unexpectedly found that the substitution rate per base-pair increases dramatically as soon as *n* exceeds ≈ 6. This effect was seen in the parent– offspring trios, the primate lineages, and in the standing variability within the human population. This observation can help explain why long stretches of guanines are avoided in coding sequences.

Most of the increase in substitution rate for G*_n_* tracks longer than *n* = 6 is due to increased G → T and G → C transversion rates, consistent with the substitutions resulting from by 8-oxo-G and other guanine lesions [14, 87–89]. In addition, we saw that in long G*_n_* tracks G → T transversions were biased towards the 5’ end of the track, in accord with *in vitro* and *in silico* studies but in stark contrast with the results for short tracks. Together, these results suggest that oxidation is especially consequential in long G*_n_* tracks, either because of an increased oxidation rate or because of a reduced effectiveness of the repair processes.

Because our aim was to assess the effect of oxidative lesions on the evolution of DNA sequences, we focused on substitutions and ignored insertions and deletions. However, like the G → T transversion rate, the rate of indels also dramatically increases with the length of a G*_n_* track [13]. It is unclear whether these phenomena have the same origin. It is possible that the indel rate is affected by oxidative stress. At the same time, the ability of G-rich regions to form G quadruplexes, which tend give rise to double double-strand breaks during DNA replication [90, 91], could also play a role. Moreover, long DNA repeats generate indels via a DNA-slippage mechanism during replication [92]; however, the frequency of polymerase slippage as well as the overall efficiency of mismatch repair system has not be found to be very dependent by sequence context [43].

In summary, the above study suggests an interesting but complex relation between *in vivo, in vitro* and *in silico* properties of DNA oxidation and the evolution of genomic sequences over diverse timescales. The mutational spectrum of short G*_n_* tracks suggests that oxidation does not play a dominant role, except in cancerous tumors. The mutational spectrum of long G*_n_* tracks showed a much increased transversion rate, but given the lack of *in vitro* or *in silico* predictions for long G*_n_* tracks, it remains uncertain what processes underly this phenomenon. Hopefully, these results will stimulate new *in vitro, in vivo,* and *in silico* studies on the rates of oxidation-caused mutations in long G*_n_* sequences.

## IV. MATERIALS AND METHODS

To obtain the *de novo* substitutions of parental genomes we combined two databases [39, 79]. This resulted in 4,933 + 11,020 = 15,953 *de novo* mutations from 78 + 250 = 328 sequenced parent–offspring trios. For each observed substitution of a G nucleotide, we calculated the length of the G_n_, track in which it is located according to the reference genome. To calculate the substituted fraction of G nucleotides at position *i* in tracks of length *n* (Fig. 1A and Fig. 2A–D), we counted the number of such substitutions and divided it by the number of G_n_, tracks in the reference genome and by the number of parent–offspring trios.

The substitutions along the primates lineages were obtained from a multiple alignment (with paralogs) [80, 81] of 7 primates (*Homo sapiens, Pan troglodytes, Gorilla gorilla, Pongo abelii, Papio anubis, Callithrix jacchus* and *Macaca mulatta*) and their derived reconstructed ancestry genome (as obtained from the same database). To estimate the mutation probability of position *i* in G tracks of length *n* (Figs 1B, 2I–L, and 4), we first identified all G nucleotides in the ancestry genome that are at position *i* in a G track of length *n.* For the set of such ancestral G nucleotides, we then counted the total number of Gs (matches) and non-Gs (mismatches) at the corresponding positions in the aligned primates. (If a primate had no alignment at a certain position, it did not contribute to either count.) The mutation probability is estimated as the the ratio of the number of matches to the sum of matches and mismatches.

To assess the variability of G_*n*_, tracks in human genomes we used the data from the 1000G project [86], containing 81377202 SNPs, obtained from 2504 individuals from 26 populations. To estimate the substituted fraction relative to the reconstructed ancestry genome for position *i* in G tracks of length *n* (Figs. 2E–I and 3), we first identified the set of all G nucleotides in the reconstructed ancestry genome that are at position *i* in a G track of length *n.* The fraction substituted by a particular nucleotide H (that is, A, C, or T), plotted in columns 2 to 4 in Fig. 3, is the fraction of nucleotides in the set that have H as an alternative allele in the database.

To analyze the mutational spectrum along G_*n*_, tracks during cancer development we exploit the Catalogue Of Somatic Mutations In Cancer [82], which contains about 15 million mutations. For each G nucleotide along the reference genome we searched the catalog for SNPs at this position. Thus, we counted the number of substitutions found at position *i* in a G track of length *n* in the reference genome. The (normalized) substituted fractions plotted in Fig. 1C are obtained by dividing these counts by the number of G tracks of length *n* in the reference genome and normalizing by the substituted fraction of G_1_ tracks (that is, 5’-HGH-3’ sequences).

